# The exon junction complex undergoes a compositional switch that alters mRNP structure and nonsense-mediated mRNA decay activity

**DOI:** 10.1101/355495

**Authors:** Justin W. Mabin, Lauren A. Woodward, Robert Patton, Zhongxia Yi, Mengxuan Jia, Vicki Wysocki, Ralf Bundschuh, Guramrit Singh

## Abstract

The exon junction complex (EJC) deposited upstream of mRNA exon junctions shapes structure, composition and fate of spliced mRNA ribonucleoprotein particles (mRNPs). To achieve this, the EJC core nucleates assembly of a dynamic shell of peripheral proteins that function in diverse post-transcriptional processes. To illuminate consequences of EJC composition change, we purified EJCs from human cells via peripheral proteins RNPS1 and CASC3. We show that EJC originates as an SR-rich mega-dalton sized RNP that contains RNPS1 but lacks CASC3. After mRNP export to the cytoplasm and before translation, the EJC undergoes a remarkable compositional and structural remodeling into an SR-devoid monomeric complex that contains CASC3. Surprisingly, RNPS1 is important for nonsense-mediated mRNA decay (NMD) in general whereas CASC3 is needed for NMD of only select mRNAs. The promotion of switch to CASC3-EJC slows down NMD. Overall, the EJC compositional switch dramatically alters mRNP structure and specifies two distinct phases of EJC-dependent NMD.

## INTRODUCTION

From the time of their birth during transcription until their eventual demise following translation, messenger RNAs (mRNA) exist decorated with proteins as mRNA-protein particles, or mRNPs (Gehring et al., 2017; Singh et al., 2015). The vast protein complement of mRNPs has been recently illuminated (Baltz et al., 2012; Castello et al., 2012; Hentze et al., 2018), and is believed to change as mRNPs progress through various life stages. However, the understanding of mechanisms and consequences of mRNP composition change remains confined to only a handful of its components. For example, mRNA export adapters are removed upon mRNP export to provide directionality to mRNP metabolic pathways, and the nuclear cap and poly(A)-tail binding proteins are exchanged for their cytoplasmic counterparts after mRNP export to promote translation (Singh et al., 2015, and references therein). When, where and how the multitude of mRNP components change during it’s lifetime, and how such changes impact mRNP function remain largely unknown.

A key component of all spliced mRNPs is the exon junction complex (EJC), which assembles during pre-mRNA splicing 24 nucleotides (nt) upstream of exon-exon junctions (Boehm and Gehring, 2016; Le Hir et al., 2016; Woodward et al., 2017). Once deposited, the EJC enhances gene expression at several post-transcriptional steps including pre-mRNA splicing, mRNA export, mRNA transport and localization, and translation. If an EJC remains bound to an mRNA downstream of a ribosome terminating translation, it stimulates rapid degradation of the mRNA via nonsense-mediated mRNA decay (NMD). The stable EJC core forms when RNA bound EIF4A3 is locked in place by RBM8A (also known as Y14) and MAGOH. It has been proposed that this trimeric complex is joined by a fourth protein CASC3 (also known as MLN51 or Barenstz) to form a stable tetrameric core (Boehm and Gehring, 2016; Hauer et al., 2016; Le Hir et al., 2016; Tange et al., 2005). However, more recent evidence suggests that CASC3 may not be present in all EJCs and may not be necessary for all EJC functions (Mao et al., 2017; Singh et al., 2012). Nonetheless, the stable EJC core is bound by a dynamic shell of peripheral EJC proteins such as pre-mRNA splicing factors (e.g. SRm160, RNPS1), mRNA export proteins (e.g. the TREX complex), translation factors (e.g. SKAR) and NMD factors (e.g. UPF3B) (Boehm and Gehring, 2016; Le Hir et al., 2016; Woodward et al., 2017). Some peripheral EJC proteins share similar functions and yet may act on different mRNAs; e.g. RNPS1 and CASC3 can both enhance NMD but may have distinct mRNA targets (Gehring et al., 2005). Thus, the peripheral EJC shell may vary between mRNPs leading to compositionally distinct mRNPs, an idea that has largely remained untested.

We previously showed that within spliced mRNPs EJCs interact with one another as well as with several SR and SR-like proteins to assemble into mega-dalton sized RNPs (Singh et al., 2012). These stable mega RNPs ensheath RNA well beyond the canonical EJC deposition site leading to RNA footprints ranging from 150-200 nt in length, suggesting that the RNA polymer within these complexes is packaged resulting in overall compact mRNP structure. Such a compact structure may facilitate mRNP navigation of the intranuclear environment, export through the nuclear pore and transport within the cytoplasm to arrive at its site of translation. Eventually, the mRNA within mRNPs must be unpacked to allow access to the translation machinery. How long do mRNPs exist in their compact states, and when, where and how are they unfurled remains yet to be understood.

Our previous observation that in human embryonic kidney (HEK293) cells CASC3 and many peripheral EJC factors are substoichiometric to the EJC core (Singh et al., 2012) spurred us to investigate variability in EJC composition. Here we use EJC purification via substoichiometric factors to reveal that EJCs first assemble into SR-rich mega-dalton sized RNPs, and then undergo a compositional switch into SR-devoid monomeric CASC3-containing EJC. The translationally repressed mRNPs, particularly those encoding ribosomal proteins, accumulate with the CASC3-bound form of the EJC. Although both the EJC forms remain active in NMD, RNPS1, a component of the SR-rich EJCs, is crucial for all tested NMD events whereas CASC3, a constituent of SR-devoid EJC, is dispensable for NMD of many transcripts. Our findings reveal a new step in the mRNP life cycle wherein EJCs, and by extension mRNPs, undergo a remarkable compositional switch that alters the mRNP structure and specifies two distinct phases of EJC dependent NMD.

## RESULTS

### RNPS1 and CASC3 associate with the EJC core in a mutually exclusive manner

We reasoned that substoichiometric EJC proteins may not interact with all EJC cores, and therefore, some of them may not interact with each other leading to variable EJC composition. To test such a prediction, we carried out immunoprecipitations (IP) of either endogenous core factor EIF4A3 or the substoichiometric EJC proteins RNPS1 and CASC3 from RNase A-treated HEK293 total cell extracts. As expected, EIF4A3 IP enriches EJC core as well as all peripheral proteins tested (Figure 1A, lane 3). In contrast, the IPs of substoichiometric factors enriched only distinct sets of proteins. CASC3 immunopurified the EJC core proteins (EIF4A3, RBM8A and MAGOH) but not the peripheral proteins ACIN1 and SAP18 (Figure 1A, lane 4; RNPS1 could not be blotted for as it co-migrates with IgG heavy chain). Conversely, following RNPS1 IP, the EJC core proteins as well as its known binding partner SAP18 (Murachelli et al., 2012; Tange et al., 2005) were enriched but CASC3 was undetected in these IPs. We saw an identical lack of co-IP between RNPS1 and CASC3 from cells that were crosslinked with formaldehyde prior to lysis (data not shown; also see below), suggesting that the lack of interaction between these substoichiometric factors does not result from their dissociation following cell lysis. We also carried out similar IPs from RNase-treated total extracts from mouse brain cortical slices, mouse embryonal carcinoma (P19) cells, and HeLa cells. Again, both RNPS1 and CASC3 efficiently co-IPed with EIF4A3 from these lysates but no co-IP was detected between RNPS1 and CASC3 (compare lanes 4 and 5 with lane 3 in Figures 1B, S1A and S1B). Therefore, we conclude that in mammalian cells RNPS1 and CASC3 exist in complex with the EJC core in a mutually exclusive manner, and term these compositionally distinct EJCs as alternate EJCs.

**Figure 1.**
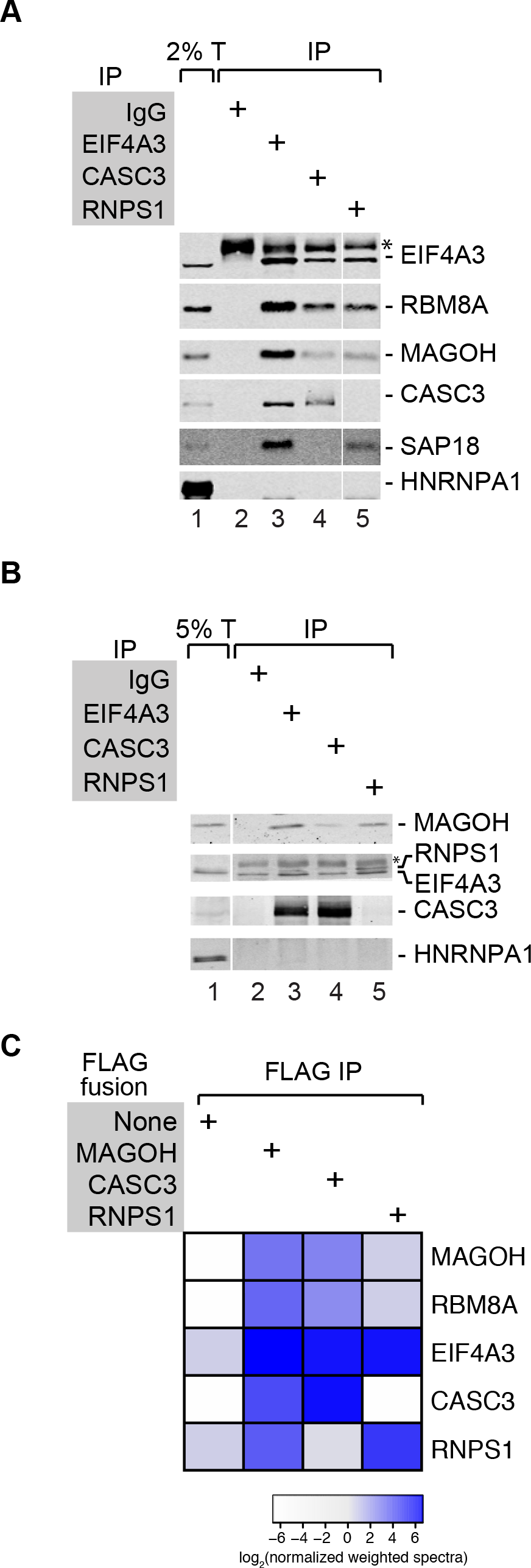
RNPS1 and CASC3 exists in mutually exclusive EJCs in mammalian cells. (A) Alternate EJCs in HEK293 cells. Western blots showing proteins on the right in total cell extract (T) (lane 1) or the immuno-precipitates (IP) (lanes 2-5) of the antibodies listed on the top. The asterisk (*) indicates signal from IgG heavy chain. (B) Alternate EJCs in mouse cortical total extracts and IPs as in (A). (C) Confirmation of mutually exclusive interactions via bottom-up proteomics. Heatmap showing signal for normalized-weighted spectra observed for the proteins on the right in the indicated FLAG IPs (top). The color scale for log_2_ transformed normalized-weighted spectral values is shown below.

### Proteomic Analysis of Alternate EJCs

To gain insights into alternate compositions of the EJC, we further characterized RNPS1 and CASC3-containing complexes using a bottom-up proteomics approach. We generated HEK293 cell lines stably expressing FLAG-tagged RNPS1 and CASC3 from a tetracycline inducible promoter where the proteins are induced at near endogenous levels (Figure S1C). FLAG IPs from RNase-treated total extracts of FLAG-RNPS1 and FLAG-CASC3 expressing cells confirmed that the tagged proteins also exist in mutually exclusive EJCs (Figure S1D). Importantly, proteomic analysis of FLAG-affinity purified alternate EJCs show almost complete lack of spectra corresponding to RNPS1 and CASC3 in the immunoprecipitates of the other alternate EJC factor (Figures 1C and S1E). In comparison, a FLAG-MAGOH IP enriches both CASC3 and RNPS1 as expected (Figures 1C, S1D and S1E).

We next compared the full complement of proteins associated with FLAG-RNPS1, FLAG-CASC3 and FLAG-MAGOH focusing on proteins >2-fold enriched in any one of the three EJC IPs as compared to the FLAG-only negative control (Table S1; see Experimental Procedures). Of the 59 proteins enriched in the two FLAG-RNPS1 biological replicates (Figure S2A), 38 are common with the 45 proteins identified in the two FLAG-MAGOH replicates (Figures S2A and S2B). Such a high degree of overlap suggests that RNPS1 containing complexes are compositionally similar to those purified via the EJC core. In comparison, among the two FLAG-CASC3 replicates the only common proteins are EIF4A3 and MAGOH (Figure S2A), which are also shared with FLAG-MAGOH (Figure S2C). Known EJC interactors such as UPF3B and PYM1 are identified in only one of the two FLAG-CASC3 replicates indicating less stable CASC3 association with these EJC factors.

A direct comparison of FLAG-RNPS1 and FLAG-CASC3 proteomes highlights their stark differences beyond the common EJC core (Figures 2A and 2B). Several SR and SR-like proteins are enriched in the RNPS1 and MAGOH proteomes but are completely absent from the CASC3 samples. Among the SR protein family, SRSF1, 6, 7 and 10 are reproducibly enriched in all RNPS1 and MAGOH samples, while all other canonical SR proteins, with the exception of non-shuttling SRSF2, are detected in at least one of the two replicates (Figure 2A, 2B, 2C and Table S1). Several SR-like proteins such as ACIN1, PNN, SRRM1 and SRRM2 are also highly enriched in MAGOH and RNPS1 IPs but are absent in CASC3 IPs (Figure 2 and Table S1). A weak but erratic signal for SRSF1 and SAP18 is seen in FLAG-CASC3 IPs (e.g. spectra for two SRSF1-derived peptides are observed in one of the FLAG-CASC3 samples; Figure 2B). However, their enrichment in CASC3-EJC is much weaker as compared to MAGOH or RNPS1 samples. Overall, these findings suggest an extensive interaction network between SR and SR-like proteins and the RNPS1-EJC but not the CASC3-EJC.

**Figure 2.**
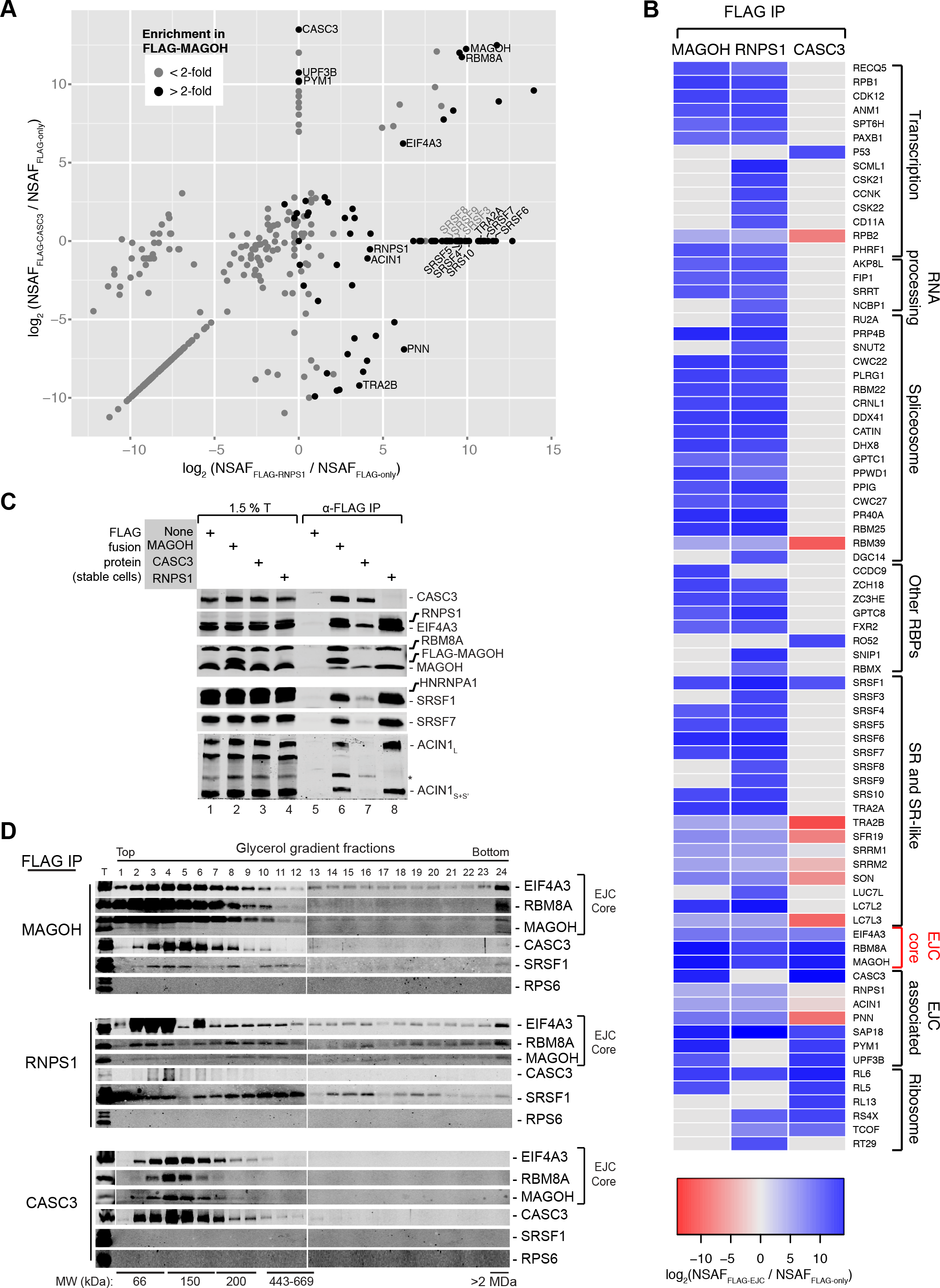
CASC3- and RNPS1-containing complexes have distinct protein composition and hydrodynamic size. (A) Comparison of alternate EJC proteomes-I. A scatter plot where log_2_ transformed values are compared for fold-enrichment of proteins in FLAG-RNPS1 IP over FLAG-only control (x-axis) to fold-enrichment of the same proteins in FLAG-CASC3 IP over FLAG-only control (y-axis). Protein quantification was performed using Normalized Spectral Abundance Factor (NSAF). Each dot represents a protein that was identified in FLAG-EJC or FLAG-only control samples by Scaffold (see experimental procedures). Black dots: proteins enriched >2-fold over control in the FLAG-Magoh IP. (B) Comparison of alternate EJC proteomes-II. A heatmap showing proteins that are >10-fold enriched in any of the FLAG-EJC IPs (indicated on the top) over FLAG-only control (proteins deemed to be contaminants were omitted). Proteins are grouped according to common functions (right). The color scale for the heatmap is at the bottom. (C) Validation of mass spec. Western blots showing proteins (right) in total extract (T) or FLAG-IPs as on the top from HEK293 cells stably expressing FLAG-tagged proteins indicated on the top left. Note that RNPS1 migrates just above EIF4A3. The asterisk (*) indicates cross-reaction with CASC3 observed with anti-ACIN1 antibody. (D) Hydrodynamic sizes of the alternate EJCs. Western blots showing proteins on the right in glycerol gradient fractions of FLAG-IPs from HEK293 cells stably expressing FLAG-MAGOH, FLAG-RNPS1, or FLAG-CASC3 (far left). The position of fractions from gradients (top) and molecular weight standards are indicated (bottom).

The proteomes of the three EJC factors can be readily classified based on protein functions in RNA biogenesis, their assembly/function within RNPs, or their protein sequence composition (Figure 2B). Many RNA biogenesis factors are specifically associated with RNPS1 (and MAGOH) but not with CASC3. These include transcription machinery components (e.g. RPB1, RPB2), transcriptional regulators (e.g. CD11A, CDK12) as well as RNA processing factors (e.g. NCBP1, FIP1). Consistent with EJC core assembly during pre-mRNA splicing, MAGOH and RNPS1 interactors also include U2, U4 and U6 snRNP components and nineteen complex subunits. None of the splicing components are seen to interact with CASC3. Considering that SR proteins also assemble onto nascent RNAs co-incident with transcription and splicing (Zhong et al., 2009), the EJC-RNPS1-SR interaction network likely originates during co-transcriptional mRNP biogenesis. In contrast, CASC3 most likely engages with the EJC only post-splicing, as previously suggested (Gehring et al., 2009a).

### Alternate EJCs differ in their higher order structure

The enrichment of SR and SR-like proteins exclusively with RNPS1 suggested that RNPS1 EJCs are likely to resemble the previously described higher order EJCs (Singh et al., 2012). Indeed, glycerol gradient fractionation of RNase-treated complexes immunopurified via FLAG-RNPS1 shows that, like EJCs purified via FLAG-MAGOH, FLAG-RNPS1 EJCs contain both lower and higher molecular weight complexes (Figure 2D). On the other hand CASC3 is mainly detected in lower molecular weight complexes purified via FLAG-MAGOH (Figure 2D, compare CASC3 signal in fractions 2-10 and 22-24). Furthermore, complexes purified via FLAG-CASC3 are exclusively comprised of lower molecular weight complexes, likely to be EJC monomers (Figure 2D). These findings are consistent with the proteomic results and suggest that compositional distinctions between the two alternate EJCs give rise to two structurally distinct complexes.

### RNPS1 and CASC3 bind RNA via the EJC core with key distinctions

We next identified the RNA binding sites for the two alternate EJC factors using RNA:protein immunoprecipitation in tandem (RIPiT) combined with high-throughput sequencing, or RIPiT-Seq (Singh et al., 2012, 2014). RIPiT-Seq entails tandem purification of two subunits of an RNP, and is well-suited to study EJC composition via sequential IP of its constant (e.g. EIF4A3, MAGOH) and variable (e.g. RNPS1, CASC3) components (Figure 3A). We carried out RIPiTs from HEK293 cells by either pulling first on FLAG-tagged alternate EJC factor followed by IP of an endogenous core factor, or vice versa. As a control, we also performed FLAG-MAGOH:EIF4A3 RIPiT-Seq. All *in vivo* EJC binding studies thus far have employed a short incubation with translation elongation inhibitor cycloheximide prior to cell lysis to limit EJC disassembly by translating ribosomes (Hauer et al., 2016; Saulière et al., 2012; Singh et al., 2012). However, to capture unperturbed, steady-state populations of RNPS1 and CASC3 bound EJCs (as in Figures 1 and 2), we performed RIPiT-Seq without translation inhibition.

**Figure 3.**
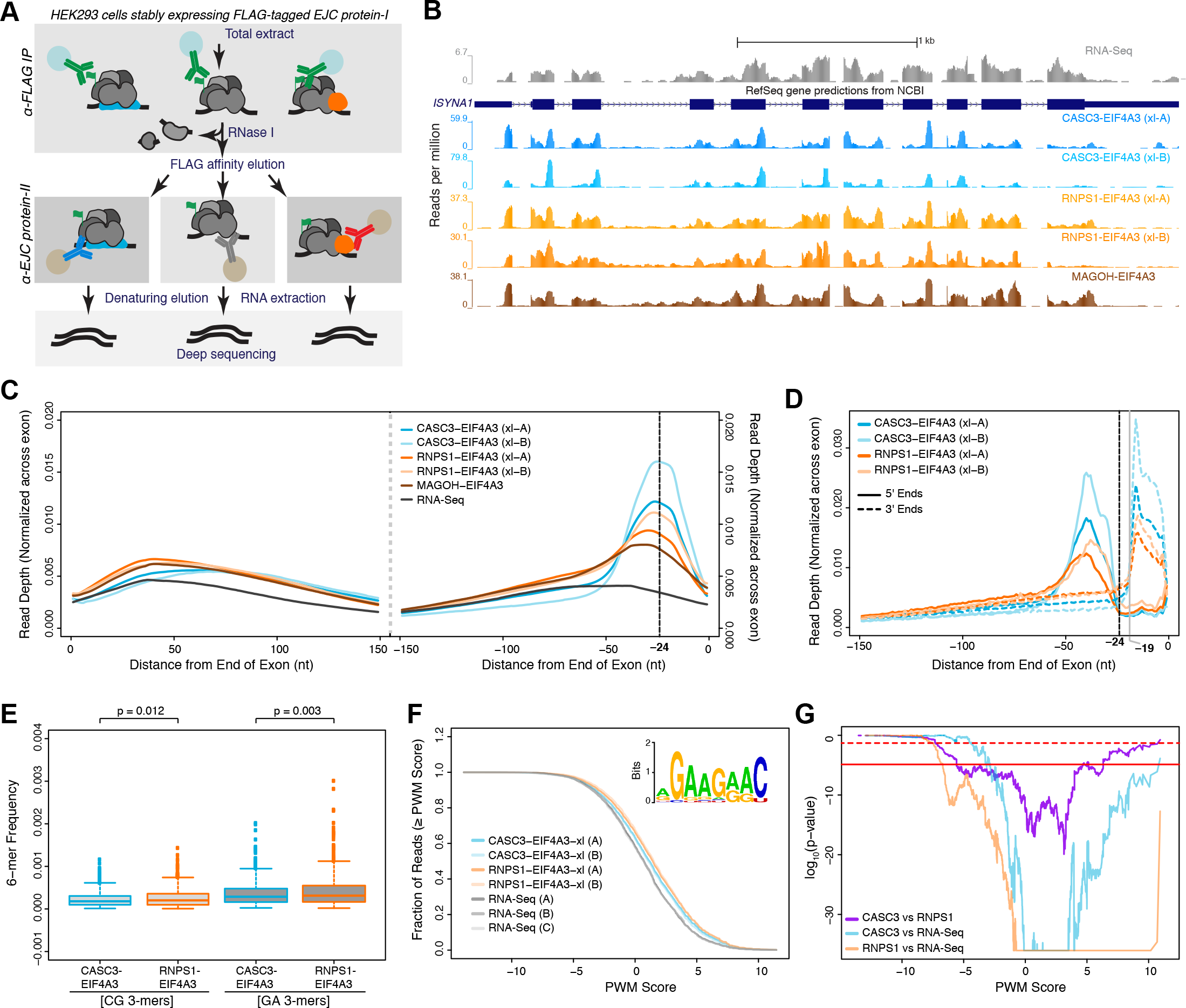
RNPS1 assembles with canonical and non-canonical EJCs but CASC3 is mainly part of canonical EJCs. (A) A schematic illustrating main steps in RNA:protein Immunoprecipitation in Tandem (RIPiT)-Seq. (B) Genome browser screen-shots comparing read coverage along the *ISYNA1* gene in RNA-Seq or RIPiT-Seq libraries (indicated on the right). The y-axis of each track in this window was normalized to millions of reads in each library. Blue rectangles: exons; thinner rectangles: untranslated regions; lines with arrows: introns. (C) Meta-exon plots showing read depth in different RIPiT-Seq or RNA-Seq libraries (indicated in the middle) in the 150 nucleotides (nt) from the exon 5’ end (left) or from the exon 3’ end (right). The vertical black dotted line marks the −24 nt position previously determined to be the site of canonical EJCs (Singh et al., 2012). (D) A composite plot of RNPS1 and CASC3 RIPiT-Seq footprint read 5’ (solid lines) and 3’ (dotted lines) ends. The canonical EJC site (−24 nt) is indicated by the vertical dotted line. Note there is some variability in the 5’ boundary of EJCs whereas the 3’ boundary is more strict (indicated by the vertical grey line at the - 19 position). (E) Box plots showing frequencies of 6-mers that contained CG 3-mers (CGG, GCG, CCG, CGC) or GA 3-mers (GGA, GAA, AGG, GAG) in CASC3 or RNPS1 RIPiT-Seq reads. Top: p-values based on Wilcoxon rank-sum test. (F) Cumulative distribution function plots showing frequency of reads in the indicated samples with the highest score for match to SRSF1 motif (inset) position weight matrix (PWM). Sample identity is in the legend on the bottom left. The SRSF1 motif shown in the inset is from (Tacke and Manley, 1995). (G) A negative binomial based assessment of significance of differences in SRSF1 motif PWM scores in (F) between the two alternate EJC footprint reads or between alternate EJC and RNA-Seq reads (legend on the bottom left). The horizontal dotted red line indicates p=0.05 whereas the horizontal solid red line indicates the Bonferroni adjusted p-value.

As expected, RIPiTs for each of the two alternate EJCs specifically purified the targeted complex along with the EJC core factors and yielded abundant RNA footprints (Figure S3A). Strand-specific RIPiT-Seq libraries from ~35-60 nucleotide footprints yielded ~5.7 to 47 million reads, of which >80% mapped uniquely to the human genome (hg38; Table S2). Genic read counts are highly correlated between RIPiTs where the order of IP of EJC core and alternate factors was reversed (Figures S3B and S3C). We found that RNPS1-EJC core interaction is susceptible to NaCl concentrations greater than 250 mM whereas CASC3-EJC core interaction persists up to 600 mM NaCl (data not shown). Therefore, to preserve labile interactions we performed alternate EJC RIPiTs from cells cross-linked with formaldehyde before cell lysis (Figure S3D). We observed a strong correlation for genic read counts between crosslinked and uncrosslinked samples (Figures S3E and S3F). The analysis presented below is from two well-correlated biological replicates of formaldehyde crosslinked RIPiT-Seq datasets of RNPS1- and CASC3-EJC (Figures S3G and S3H).

Consistent with EJC deposition on spliced RNAs, alternate EJC footprints are enriched in exonic sequences from multi-exon genes (Figure S3I). Along individual exons, footprints of both alternate EJCs mainly occur close to exon 3’ ends (Figure 3B). Indeed, a meta-exon analysis shows that the alternate EJC factors bind mainly to the canonical EJC binding site 24 nt upstream of exon junctions (Figure 3C) (Hauer et al., 2016; Saulière et al., 2012; Singh et al., 2012). Notably, the alternate EJC footprint enrichment at the canonical EJC site is irrespective of whether the alternate EJC factors are IPed first or second during the RIPiT procedure (Figure S3J). Thus, both RNPS1 and CASC3 mainly bind to RNA via EJCs at the canonical site. The location of the 5’-and 3’-ends of the alternate EJC footprint reads shows that both alternate EJCs have a similar footprint on RNA as each blocks positions −26 to −19 nt from RNase I cleavage (Figure 3D). Our findings are consistent with previous work that showed that CASC3 mainly binds to the canonical EJC site (Hauer et al., 2016). However, in contrast to this study, RNPS1 binding to canonical EJC sites is readily apparent (Figures 3B, 3C and 3D), which could reflect differences in the UV-crosslinking dependent CLIP-Seq approach used by Hauer *et al.* and the photo-crosslinking independent RIPiT-Seq. Altogether, our results suggest that both RNPS1 and CASC3 exist in complex with EJC at its canonical site.

While read densities for both alternate EJC factors are highest at the canonical EJC site, 47-62% of reads map outside of the canonical EJC site similar to previous estimates (Saulière et al., 2012; Singh et al., 2012). Non-canonical footprints are somewhat more prevalent in RNPS1-EJC as compared to CASC3-EJC (Figure S3K). As RNPS1-EJC is intimately associated with SR and SR-like proteins (Figure 2A and B), a k-mer enrichment analysis revealed a modest but significant enrichment of GA-rich 6-mers in RNPS1 over CASC3 footprints (Figures 3E). Such purine-rich sequences occur in binding sites of several SR proteins (SRSF1, SRSF4, Tra2a and b) (Änkö et al., 2012; Pandit et al., 2013; Tacke et al., 1998). There is also a small but significant enrichment of sequence motifs recognized by SRSF1 and SRSF9 in RNPS1 footprints as compared to CASC3 footprints or RNA-Seq reads (Figures 3F, 3G, S3L and S3M). We conclude that within spliced RNPs, RNPS1-containing EJC is engaged with SR and SR-like proteins, and other RNA binding proteins, which leads to co-enrichment of RNA binding sites of these proteins during RIPiT via RNPS1. Intriguingly, CG-rich 3-mers are the highest enriched k-mers in footprints of both alternate EJCs as compared to RNA-Seq (data not shown). The CG-rich 6-mers are also somewhat more enriched in RNPS1-EJC (Figure 3E). A CG-rich sequence was reported as an *in vitro* RBM8A binding motif indicating a yet to be determined relationship between the EJC core and CG-rich sequences (Ray et al., 2013).

### RNPS1 and CASC3 RNA occupancy changes with mRNP subcellular location

Surprisingly, despite the mutually exclusive association of RNPS1 and CASC3 with the EJC core (Figures 1 and 2), the two proteins are often detected on the same sites on RNA leading to their similar apparent occupancy on individual exons as well as entire transcripts (Figures 3B, S4A and S4B). These results suggest that the two alternate EJC factors bind to two distinct pools of the same RNAs. To further reveal RNA binding patterns of the alternate EJC factors, we first identified exons differentially enriched in one or the other factor. Remarkably, if two or more exons of the same gene are differentially enriched in RNPS1 or CASC3 footprints, these exons are almost always enriched in the same alternate EJC factor (Figure 4A, p < 0.00001). This extremely tight linkage between EJC compositions of different exons of the same gene suggests that RNPS1 and CASC3 binding to EJC is determined at the level of the entire transcript and not of the individual exon. At any given time a transcript with multiple EJCs is likely to be homogeneously associated with either one or the other alternate factor. In support of this scenario, very weak RNA-dependent interaction is detected between the two alternate EJC factors (Figure S4C).

**Figure 4.**
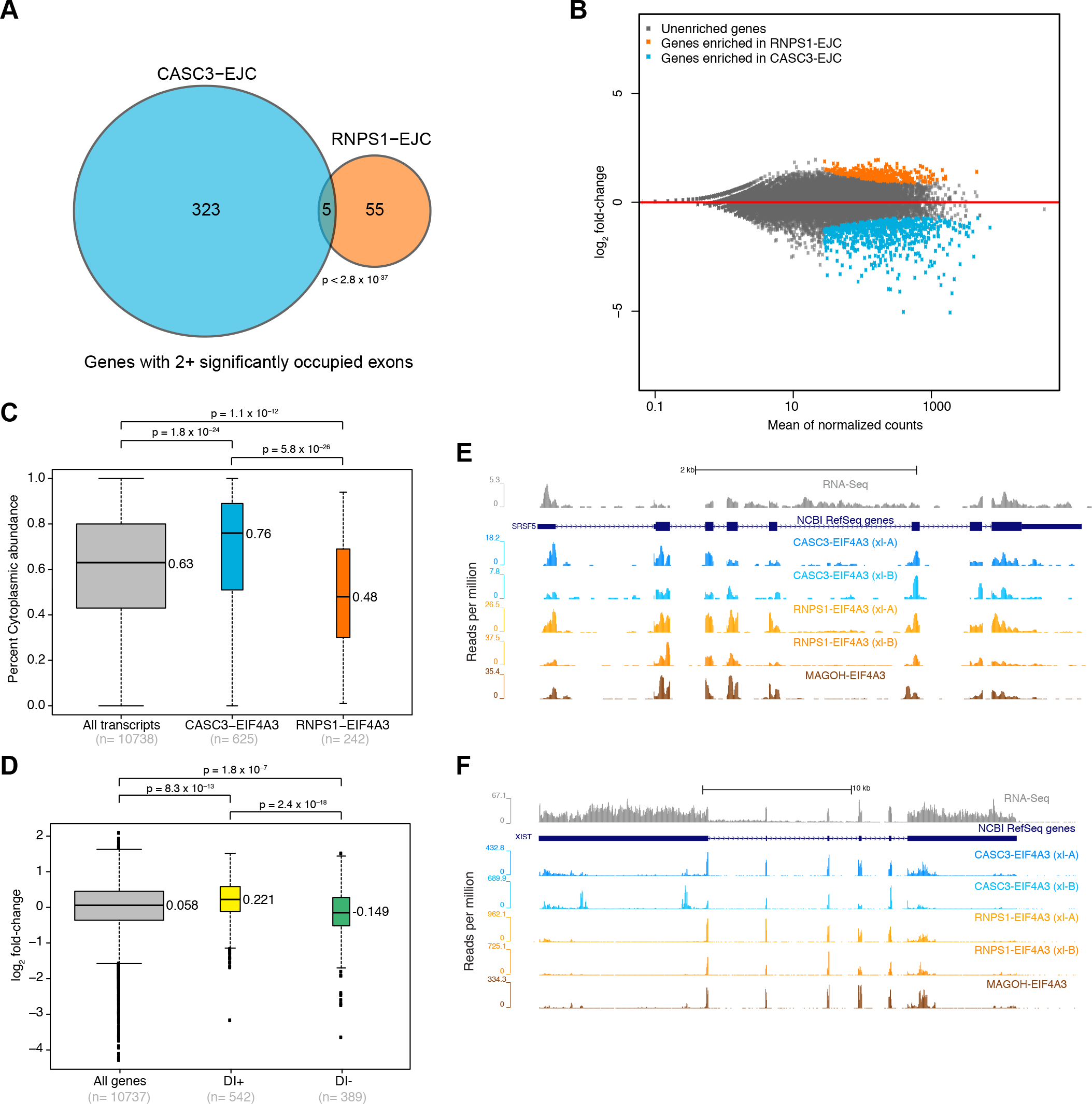
RNPS1 and CASC3 occupy same RNAs and sites but in different subcellular compartments. (A) A venn-diagram showing genes with at least two differentially enriched exons, for which all differentially enriched exons are preferentially bound to CASC3-EJC (blue region), RNPS1-EJC (orange region), or genes with at least one exon is enriched in CASC3-EJC and one in RNPS1-EJC (overlap region). (p < 2.8 × 10^−37^). p-value shown was obtained as the probability of observing an overlap of 5 or fewer genes using a binomial distribution approximating each gene as containing two exons. (B) An MA-plot showing fold-change in RNPS1-EJC versus CASC3-EJC footprint reads (y-axis) against expression levels (x-axis). Each dot represents a canonical transcript for each known gene in GRChg38 from UCSC “knownCanonical” splice variant table. Transcripts determined to be differentially enriched (p-adjusted<0.05) in RNPS1-EJC (orange) and CASC3-EJC (blue) are indicated. (C) Box plots showing distribution of percent cytoplasmic levels (y-axis) for all (grey), CASC3-EJC enriched (blue), and RNPS1-EJC enriched (orange) transcripts. The median values are given to the right of each box plot. p-values on the top are based on Wilcoxon rank sum test. Bottom: The number of transcripts in each group. (D) Box plots showing distribution of fold-change values (y-axis) for RNPS1-EJC versus CASC3-EJC footprint read counts for all (grey), detained intron-containing (DI+; yellow) or detained intron-lacking (DI-; green) transcripts. (+ve values: enriched in RNPS1-EJC; -ve values: enriched in CASC3-EJC). Top: p-values (Wilcoxon rank sum test). Bottom: the number of transcripts in each group. (E) Genome browser screen-shots showing read coverage along the *SRSF5* gene in RNA-Seq (top) or RIPiT-Seq libraries (labeled on right). Blue rectangles: exons; thinner rectangles: untranslated regions; lines with arrows: introns. Note increased RNA-Seq reads in introns 4 and 5. (F) Genome browser screen-shot as in (E) for the *XIST* locus.

At steady state, RNPS1 is mainly localized to the nucleus whereas CASC3 is predominantly cytoplasmic although both proteins shuttle between the two compartments (Daguenet et al., 2012; Degot et al., 2002; Lykke-Andersen et al., 2001). We reasoned that different concentration of alternate EJC factors in the two subcellular compartments may mirror their EJC association and RNA occupancy. To test this possibility, we identified subsets of transcripts that are preferentially enriched in RNPS1-(242 transcripts) or CASC3-EJC (625 transcripts; Figure 4B), and compared their nuclear:cytoplasmic ratios based on estimates of subcellular RNA distribution in HEK293 cells (Neve et al., 2016). Indeed, the transcripts enriched in CASC3-EJC strongly localize to the cytoplasm (median % localized in cytoplasm = 76), whereas those preferentially bound to RNPS1 show more even distribution with a slight bias for nuclear localization (median % localized in cytoplasm = 48; Figures 4C and S4D). Thus, subcellular localization of spliced RNAs and EJC factors are important determinants of EJC composition.

We surmised that kinetics of mRNA maturation and in turn nuclear export will directly impact EJC composition. To test this idea, we compared alternate EJC occupancy of a group of genes whose nuclear poly(A)-tailed transcripts contain introns that undergo splicing much more slowly as compared to other introns in the same transcript (Boutz et al., 2015). Transcripts containing these detained introns (DI) are restricted to the nucleus in a mostly pre-processed state until DIs are spliced, and are therefore expected to have slower mRNA export rates. Of the genes that Boutz *et al.* found to contain DI in four human cell lines, 542 also contained DIs in HEK293 cells whereas a set of 389 transcripts do not contain DI in any human cell lines including HEK293 cells (data not shown). We find that DI-containing transcripts are significantly more enriched in RNPS1-EJC (p=8.3×10^−13^, Figures 4D and 4E) while DI lacking transcripts show an enrichment in CASC3-EJC (p=1.8×10^−7^). These results support our conclusion that EJC (and mRNP) compositional switch mainly occurs as they mature and enter the cytoplasm, and that the rate of mRNA progression to the cytoplasm alters the switch.

Despite a strong CASC3 enrichment on cytoplasmic RNAs, a quarter of all preferentially CASC3-bound RNAs are more nuclear (Figure 4C). Consistent with CASC3 shuttling into the nucleus (Daguenet et al., 2012; Degot et al., 2002), its footprints are abundantly detected on *XIST* RNA and several other spliced non-coding RNAs restricted or enriched in the nucleus (Figures 4F and S4D). Further, in HEK293 nuclear as well as cytoplasmic extracts both CASC3 and RNPS1 are found to assemble into EJCs (Figures S4E and S4F). These data suggest that while the bulk of RNPs may undergo the switch in EJC composition in the cytoplasm, some RNPs can undergo this change in the nucleus itself, possibly as a function of RNP half-life or due to a more active nuclear role of CASC3 in RNP biogenesis and function.

### Translation and mRNA decay kinetics impacts alternate EJC bound mRNA pools

As EJCs are disassembled during translation (Dostie and Dreyfuss, 2002; Gehring et al., 2009b), abundant RNPS1 and CASC3 footprints at EJC deposition sites suggest that the bulk of mRNPs undergo the compositional switch before translation. To test how translation impacts occupancy of both alternate EJCs, and if their occupancy is influenced by rate at which mRNAs enter translation pool, we obtained RNPS1- and CASC3-EJC footprints from cells treated with cycloheximide (CHX), and compared them to alternate EJC footprints from untreated cells. When mRNAs bound to each alternate EJC are compared across the two conditions, CASC3 RNA occupancy shows a dramatic change (Figure 5A). In contrast, while changes in RNPS1-bound RNAs across two conditions shows the same trend as CASC3 (R^2^=0.27, Figure 5A), the change is much less dramatic with RNPS1 occupancy changing significantly only on a handful of transcripts. These observations further support our conclusion that the RNPS1-bound form of the EJC precedes the CASC3-bound form. They also indicate that translation inhibition does not interfere with the compositional switch from RNPS1 to CASC3 but leads to accumulation of CASC3-bound mRNPs.

**Figure 5.**
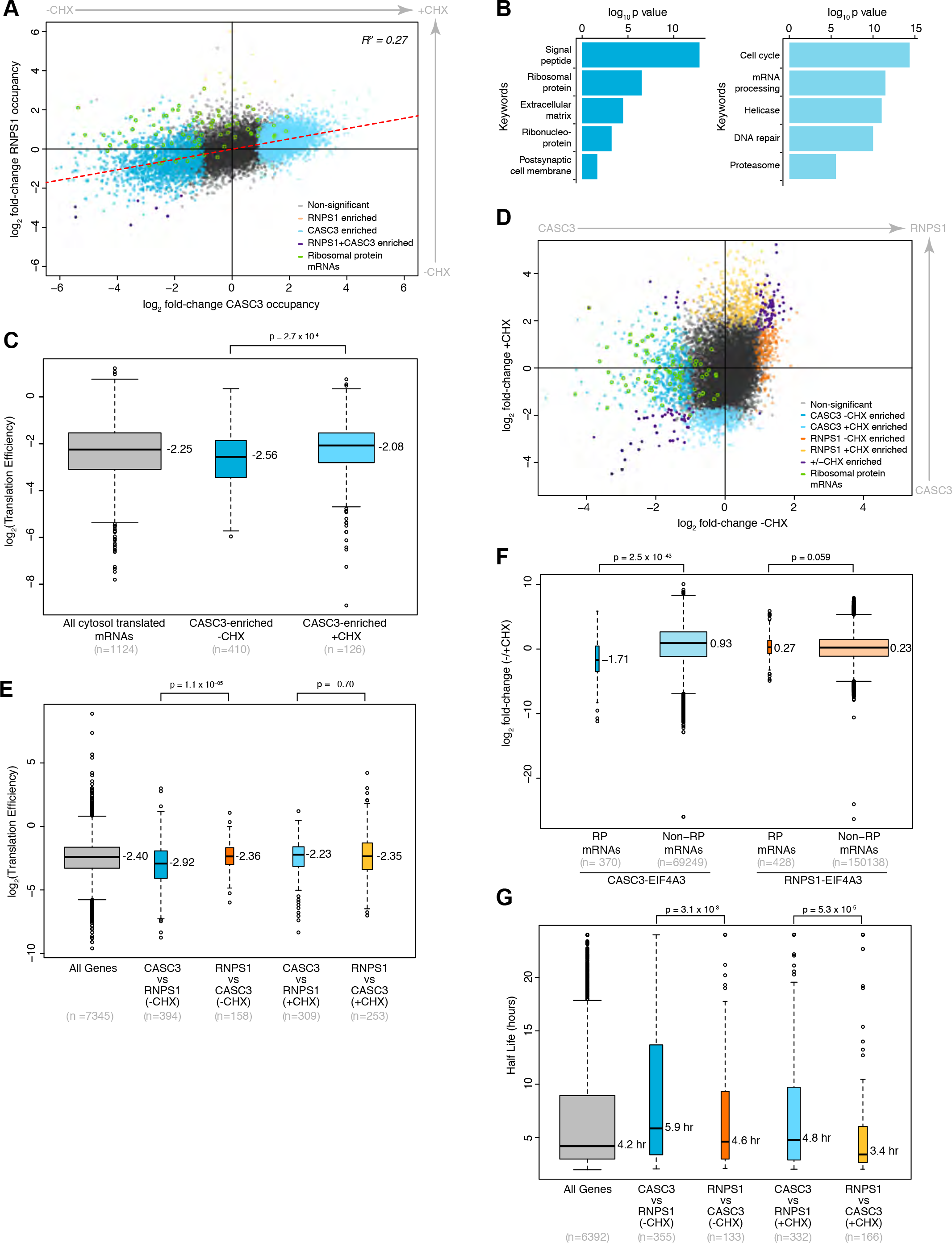
Translation and mRNA decay kinetics impacts alternate EJC occupancy. (A) A scatter plot showing fold-change in CASC3-EJC occupancy in absence versus presence of cycloheximide (CHX; x-axis) and fold-change in RNPS1-EJC in absence versus presence of CHX (y-axis). Each dot represents a canonical transcript for each known gene in GRChg38, and is colored as indicated in the legend (bottom right). The dots representing ribosomal protein genes are highlighted by a green outline. The dotted red line shows a linear fit and the coefficient of determination (R^2^) is on the top left corner. (B) Top 5 GO term keywords and their log_10_ transformed p-values of enrichment in CASC3-EJC enriched transcripts (from A) in the absence (left) or presence of CHX (right). (C) Box plots showing distribution of translation efficiency estimates (y-axis) of transcript groups on y-axis. The median values are given to the right of each box plot. Top: p-values (Wilcoxon rank sum test). Bottom: the number of transcripts in each group. (D) Scatter plot as in (A) comparing fold-change in CASC3-EJC versus RNPS1-EJC footprint read counts in -CHX (x-axis) and +CHX (y-axis) conditions. (E) Box plots as in (C) showing distribution of translation efficiency estimates of transcript groups from (D) as indicated on the bottom. (F) Comparison of fold-change in CASC3-EJC or RNPS1-EJC footprint reads at canonical EJC sites (as defined in the methods section) from ribosomal protein (RP)-coding mRNAs or non-ribosomal (non-RP)-protein coding mRNAs. Top: p-values (Wilcoxon rank sum test). (G) Box plots as in (C) and (E) above comparing mRNA half-life of transcript groups from (D) as indicated on the bottom.

It is expected that poorly translated or translationally repressed mRNAs will be enriched in CASC3-EJC under normal conditions, and more efficiently translated mRNAs will be differentially enriched upon translation inhibition. To test this hypothesis, we obtained mRNA translation efficiency (TE) estimates based on ribosome footprint counts for each transcript normalized to its abundance from a human colorectal cancer cell line (Kiss et al., 2017). A comparison of the CASC3-EJC enriched transcripts from the two conditions however showed only a minor difference in their median TE (Figure S5A). A search for functionally related genes in the two sets revealed that each contains diverse groups (Figure 5B). Among the transcripts differentially bound to CASC3-EJC under normal conditions, the largest and most significantly enriched group encodes signal-peptide bearing secretory/membrane bound proteins (Figure 5B), which has a significantly higher TE as compared to all transcripts (Figure S5B). We reason that despite the higher TE of the “secrotome” (all secreted/membrane proteins as classified in (Jan et al., 2014)), transcripts with signal peptide may be enriched in CASC3-EJC because binding of the hydrophobic signal peptide to the signal recognition particle halts translation, which resumes only when the ribosome engages with the endoplasmic reticulum (ER) (Walter and Blobel, 1981). Presumably, the time before translation resumption on the ER allows capture of mRNPs where EJC composition has switched but the complex has not yet been disassembled by translation. When we excluded the secretome and considered only those transcripts that are translated in the cytosol (as defined in (Chen et al., 2011)), a comparison of TE of CASC3-EJC enriched transcripts under translation conducive and inhibitory conditions confirmed our initial hypothesis. As seen in Figure 5C, transcripts bound to CASC3-EJC in the absence of CHX have significantly lower median TE (−2.56) as compared to median TE of transcripts bound to CASC3-EJC in the presence of CHX (−2.08, p=2.7 × 10^−4^).

Another functional group enriched in CASC3-EJC under normal conditions comprises the ribosomal protein (RP)-coding mRNAs (Figure 5B). Strikingly, transcripts encoding ~50% of all cytosolic ribosomal subunit proteins as well as 13 mitochondrial ribosome subunits are among this set (Figure 5A). Although RP mRNAs are among the most highly translated in the cell, a sizeable fraction of RP mRNAs are known to exist in a dormant untranslated state that can enter the translation pool upon demand (Geyer et al., 1982; Meyuhas and Kahan, 2015; Patursky-Polischuk et al., 2009). Consistently, RP mRNAs have significantly low TE in human and mouse cells (Figures S5B and S5C). Furthermore, when transcripts differentially bound to RNPS1- versus CASC3-EJC are directly compared in normally translating cells, RP mRNAs are specifically enriched among CASC3-EJC bound transcripts (Figure 5D). Therefore, RP mRNPs, and perhaps other translationally repressed mRNPs, that linger in the untranslated state in the cytoplasm switch to and persist in the CASC3-bound form of the EJC. Such a possibility is further supported by significantly lower TE of CASC3-EJC enriched RNAs as compared to RNPS1-EJC bound transcripts under normal conditions (Figure 5E). Further, when mRNPs are forced to persist in an untranslated state upon CHX treatment, cytosol translated mRNAs show increased CASC3 occupancy whereas their RNPS1 occupancy is not affected (Figure S5D).

As previously reported (Hauer et al., 2016), RP mRNAs are depleted of CASC3-EJC upon translation inhibition (Figures 5A and 5D). Upon cycloheximide treatment, CASC3 occupancy is significantly reduced at canonical EJC sites of RP mRNAs as compared to non-RP mRNAs, which show an increase in CASC3 occupancy (Figure 5F, p=2.5×10^−43^). In comparison, RNPS1 occupancy on all transcripts modestly increases upon translation inhibition (Figure 5F, p=0.059). A paradoxical possibility is that the untranslated reserves of RP mRNAs may undergo translation when the cellular pool of free ribosomes is dramatically reduced upon CHX-mediated arrest of translating ribosomes on mRNAs. Intriguingly, a recent study in fission yeast found a similar contradictory increase in ribosome footprint densities on RP mRNAs upon cycloheximide treatment (Duncan and Mata, 2017).

The alternate EJC occupancy landscape is also impacted by mRNA decay kinetics. CASC3-EJC enriched RNAs have longer half-lives as compared to RNPS1-EJC enriched transcripts under both translation conducive (median t_1/2_=4.6 hr vs. 5.9 hr, p=3.1×10^−3^) and inhibitory conditions (median t_1/2_=3.4 hr vs. 4.8 hr, p=5.3×10^−5^, Figure 5G). Notably, RNAs enriched in both alternate EJCs upon CHX treatment have lower half-lives as compared to the corresponding cohorts enriched from normal conditions. Thus, EJC detection is enhanced on transcripts that are stabilized after CHX treatment. Such a conclusion is further supported by enrichment of functionally related groups of genes known to encode unstable transcripts (e.g. cell cycle, mRNA processing and DNA damage, (Schwanhäusser et al., 2011)) in CASC3-EJC upon translation inhibition (Figure 5B).

### RNPS1 is required for efficient NMD of all transcripts whereas CASC3 is dispensable for many

Our data suggests that mRNPs arrive in the cytoplasm with RNPS1 and SR protein-containing EJCs, which are remodeled into CACS3-containing EJCs. We wanted to determine if alternate EJCs were equally important for NMD and/or if they have distinct targets as previously suggested (Gehring et al., 2005). In HEK293 cells with ~80% of RNPS1 mRNA and proteins depleted, a majority of endogenous NMD targets tested are significantly upregulated (Figures 6A and S6A). Surprisingly, however, in cells with ~85% of CASC3 depleted, NMD targets are largely unaffected; some others are only modestly upregulated, and only one out of thirteen is significantly upregulated. Notably, the RNAs that were upregulated upon CASC3 depletion are even more upregulated upon RNPS1 depletion. Most tested RNAs are significantly upregulated upon depletion of EIF4A3 and the central NMD factor UPF1 (Figure 6A and data not shown). These data suggest that RNPS1 function is required for efficient NMD of most endogenous NMD substrates whereas CASC3 may be needed for NMD of only select RNAs.

**Figure 6.**
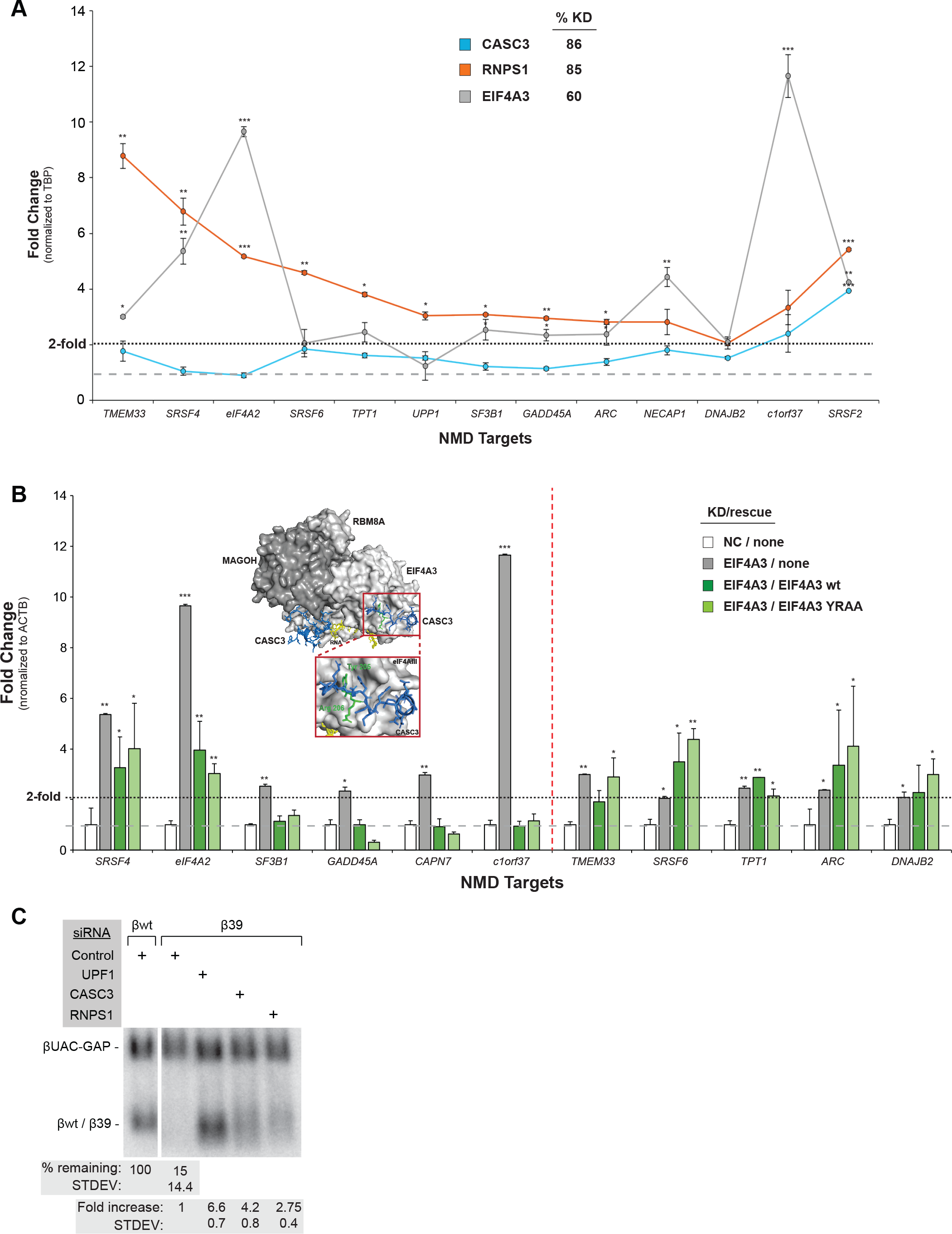
RNPS1 is required whereas CASC3 is largely dispensable for efficient NMD. (A) Fold change as measured by real-time PCR in abundance of endogenous NMD-targeted transcripts (bottom) in HEK293 cells depleted of core or alternate EJC factors. (EIF4A3 knockdown: 48 hr, alternate EJC factor knockdown: 96 hr). Shown are the average values normalized to TBP levels from three biological replicates ± standard error of means (SEM). Asterisks denote statistically significant differences: * p<0.05, ** p<0.01, *** p<0.001 (Welch’s t-test). Percent knockdown of each protein in a representative experiment is in the legend (see Figure S6). (B) Real-time PCR analysis as in (A) from HEK293 cells depleted of EIF4A3. Either wild-type EIF4A3 or a mutant lacking CASC3 interaction were exogenously expressed in EIF4A3 knockdown cells as indicated in the legend on the top right. Inset: An EJC core crystal structure (PDB ID: 2J0S) showing the EIF4A3 residues that interact with CASC3 (enlarged below). The vertical red dotted line divides the NMD targets into two groups based on the degree of NMD restoration. Northern blots showing levels of wild-type (βwt, lane 1) or PTC-containing (codon 39; β39; lanes 2-5) β-globin mRNA and a longer internal control β-globin (βUAC-GAP) mRNA from HeLa Tet-off cells treated with siRNAs indicated on the top. Tables below indicate percentage of normalized β39 mRNA as compared to normalized βwt mRNA (top) or fold-increase in β39 mRNA upon knockdown as compared to control.

It is possible that the residual amount of CASC3 after its siRNA mediated depletion is sufficient to support NMD. To test this further, we depleted endogenous EIF4A3 protein to ~50% of its normal levels and supplemented cells with either a WT EIF4A3 (WT) or an EIF4A3 mutant (YRAA: Y205A, R206A) with much reduced CASC3 binding (Figures 6B, S6B, S6C, S6D, (Andersen et al., 2006; Ballut et al., 2005; Bono et al., 2006). The EIF4A3 knockdown results in >2-fold upregulation of all NMD targets tested (Figure 6B). We find that complementation of EIF4A3 knockdown cells with exogenous FLAG-EIF4A3 or FLAG-EIF4A3 YRAA proteins leads to nearly identical effect on NMD restoration (Figure 6B). Based on the degree of rescue, the NMD targets tested can be divided into two groups. The first group (to the left of vertical red dotted line in Figure 6B) shows a strong rescue of NMD upon complementation with both the wild-type or the mutant EIF4A3 proteins. Therefore, this group of transcripts can undergo NMD largely independent of CASC3. In the second group (to the right of vertical red dotted line in Figure 6B), neither the wild-type nor the mutant proteins can rescue NMD defect caused by the depletion of endogenous EIF4A3 protein. We noted that, unlike the first group (with exception of C1orf37), the second group is somewhat more sensitive to depletion of CASC3 (Figure 6A). This subset of mRNAs may have a more complex dependence on EIF4A3 levels.

In contrast to most endogenous NMD substrates, we found that knockdown of both RNPS1 and CASC3 in HeLa cells leads to accumulation of a well-known β-globin mRNA with a premature stop codon at codon 39 (β39, Figures 6C and S6E). Consistently, recent genome-wide screens have identified CASC3 as an effector of NMD of exogenous reporters (Alexandrov et al., 2017; Baird et al., 2018). Overall, our results show that, in human cells, several NMD targets can undergo CASC3 independent NMD whereas almost all tested transcripts depend on RNPS1 for their efficient NMD. On the other hand, exogenously (over) expressed NMD substrates may depend on both sequential EJC compositions for their efficient NMD.

### Increased CASC3 levels slow down NMD

We next tested if overexpression of RNPS1 or CASC3 can tilt EJC composition toward one of the two alternate EJCs, and if such a change can impact NMD. CASC3 overexpression in HEK293 cells leads to several fold increase in CASC3 co-IP with EIF4A3 (Figures 7A, 7B) and RBM8A (Figures S7A and S7B), although no concomitant decrease is seen in RNPS1 co-precipitation with EJC core proteins. Consistent with this, manifold overexpression of RNPS1 did not cause any detectable change in levels of the two alternate factors in the EJC core IPs (Figures 7A, 7B and S7A and S7B). These results rule out a simple, direct competition between the two proteins for EJC core interaction. We next tested if increased CASC3 association with EJCs under elevated CASC3 levels affects NMD. Surprisingly, more than half of tested endogenous NMD targets are significantly upregulated upon CASC3 overexpression (Figure 7C). Similarly, the β39 mRNA exogenously expressed in HeLa cells is also modestly stabilized upon CASC3 overexpression (Figure 7D and S7C). As previously reported (Viegas et al., 2007), RNPS1 overexpression further downregulates β39 mRNA in HeLa cells (Figures S7C and S7D), indicating that NMD of this RNA occurs more efficiently when it is associated with early acting SR-rich EJC.

**Figure 7.**
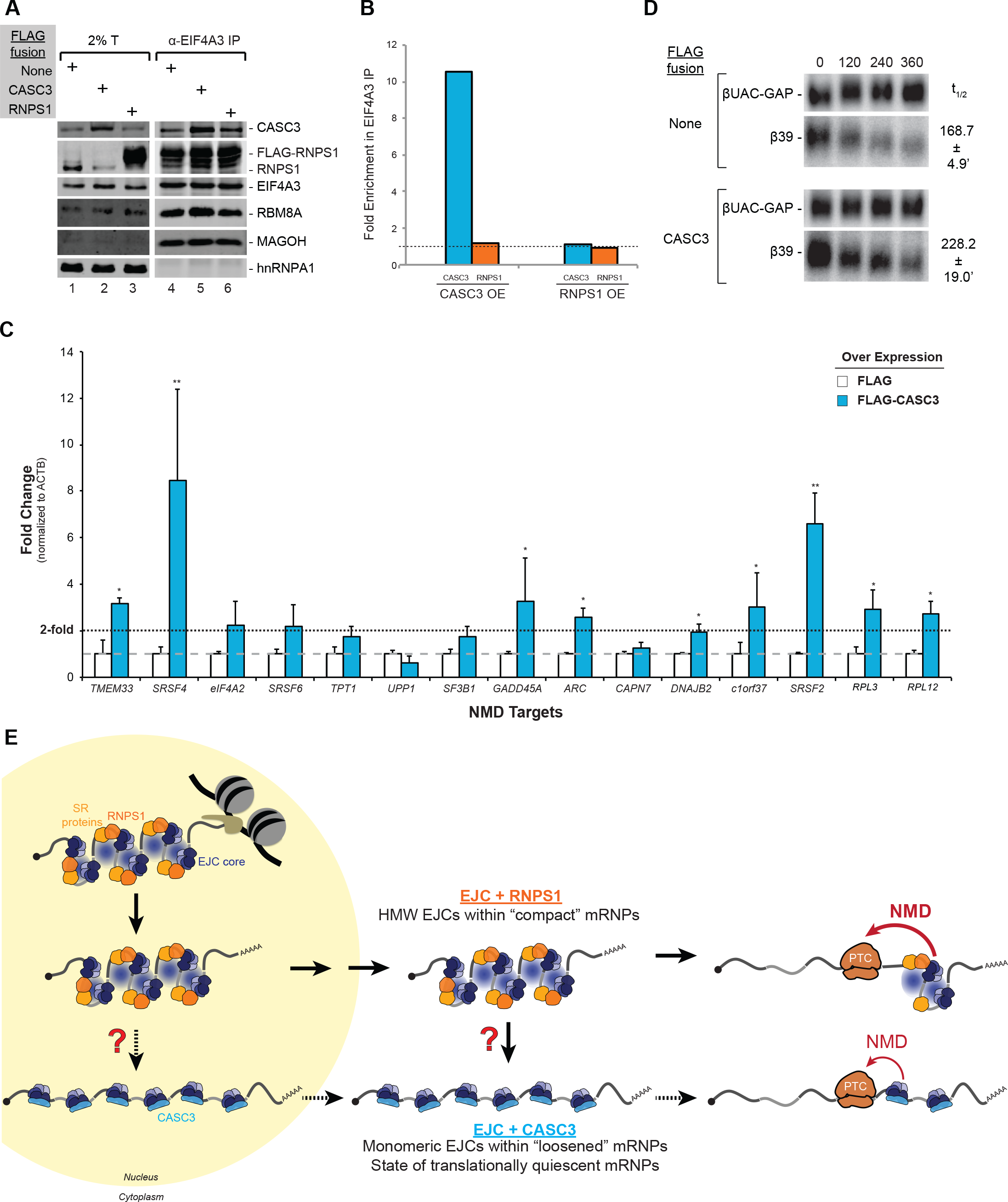
Promotion of switch to late-acting CASC3-EJC dampens NMD activity. (A) Western blots showing proteins on the right in total extract (T, lanes 1-3) or EIF4AIII IPs (lanes 4-6) from HEK293 cells overexpressing FLAG-fusion proteins at top left. (B) Histogram showing the fold enrichment of alternate EJC factors in EIF4A3 IPs from (A). Overexpressed (OE) alternate EJC factors are at the bottom and CASC3 (blue) and RNPS1 (orange) levels in each OE sample (lane 5 or 6) were compared to the control IP in lane 4 (grey, set to 1). (C) Fold change in abundance of NMD-targeted endogenous transcripts (bottom) in HEK293 cells exogenously overexpressing FLAG or FLAG-CASC3. Shown are the average values normalized to ACTB levels from three biological replicates ± SEM. Asterisks denote statistically significant differences: * p<0.05, ** p<0.01, *** p<0.001 (Welch’s t-test). (D) Northern blots showing decay of Tetracycline-inducible β39 mRNA as compared to constitutively expressed βUAC-GAP mRNA in HeLa Tet-off cells overexpressing FLAG-tagged proteins indicated on the left. Time after Tet-mediated transcriptional shut-off of β39 mRNA is indicated on the top. On the right is β39 mRNA half-life (t_1/2_) from each condition (average of three biological replicates ± standard deviation). (E) A model depicting switch in EJC composition and its effect on mRNP structure and NMD activity. Key EJC proteins are indicated. Oval with radial blue fill in the high molecular weight (HMW) EJCs represent unknown interactions that mediate EJC multimerization. Grey and black lines: exons.

## DISCUSSION

The EJC is a cornerstone of all spliced mRNPs, and interacts with upwards of 50 proteins to connect the bound RNA to a wide variety of post-transcriptional events. The EJC is thus widely presumed to be “dynamic”. By purifying EJC via key peripheral proteins, we demonstrate that a remarkable binary switch occurs in EJC’s complement of bound proteins. Such an EJC composition change has important implications for mRNP structure and function including mRNA regulation via NMD (Figure 7E).

### EJC composition and mRNP structure

Our findings suggests that when EJCs first assemble during co-transcriptional splicing, the core complex consisting of EIF4A3, RBM8A and MAGOH engages with SR proteins and SR-like factors including RNPS1 (Figure 2). Within these complexes, RNPS1 is likely bound to both the EJC core as well as to the SR and SR-like proteins bound to their cognate binding sites on the RNA (Figure 3). This network of interactions bridges adjacent and distant stretches of mRNA, winding the mRNA up into a higher-order structure, which is characteristic of RNPs purified from human cells via the EJC core proteins or RNPS1 (Figure 2, (Singh et al., 2012)). Such higher-order mRNA packaging can create a compact RNP particle for its efficient intracellular transport. It is possible that assembly of these higher-order structures occur via multiple weak interactions among low complexity sequences (LCS) within EJC bound SR and SR-like proteins (Haynes and Iakoucheva, 2006; Kwon et al., 2014). It also remains to be tested if RNPS1, which possesses an SR-rich LCS, acts as a bridge between the EJC core and more distantly bound SR proteins. Our data also indicates that on a majority of mRNPs, these SR-rich and RNPS1-containing higher-order EJCs persist throughout their nuclear lifetime (Figure 4). When mRNPs arrive in the cytoplasm, the SR and SR-like proteins are evicted from all EJCs of an mRNP and the EJC is joined by CASC3 (Figure 4). It remains to be seen if CASC3 incorporation into an EJC causes its remodeling. Alternatively, active process(es) such as RNP modification via SR protein phosphorylation by cytoplasmic SR protein kinases (Zhou and Fu, 2013) or RNP remodeling by ATPases may precede CASC3 binding to EJC (Lee and Lykke-Andersen, 2013). What is clear is that CASC3-bound EJCs lose their higher order structure and exist as monomeric complexes at the sites where EJC cores were co-transcriptionally deposited. Thus, the switch in EJC composition from RNPS1 and SR-rich complexes to CASC3-bound complexes leads to a striking alteration in higher-order EJC, and possibly, mRNP structure (Figure 7E). The CASC3 bound form of the EJC following compositional switch is the main target of translation dependent disassembly although RNPS1-EJC may undergo some translation-dependent disassembly (Figure 5).

### CASC3-EJC and pre-translation mRNPs

Our findings support the emerging view that CASC3 is not an obligate component of all EJC cores. A population of assembled EJCs, especially those early in their lifetime, completely lack CASC3 (Figures 1 and 2). Such a view of partial CASC3 dispensability for EJC structure and function is in agreement with findings from *Drosophila* where the assembled trimeric EJC core as well as RNPS1 and its partner ACIN1 are required for splicing of long or sub-optimal introns whereas CASC3 is not (Hayashi et al., 2014; Malone et al., 2014; Roignant and Treisman, 2010). A recent report that lack of CASC3 during mouse embryonic brain development results in phenotypes distinct from those caused by similar mutations of the other three core proteins also supports nonoverlapping functions of CASC3 and the other core factors (Mao et al., 2017). We note that in the recently reported human spliceosome C* structure, CASC3 is bound to the trimeric core (Zhang et al., 2017). As the spliceosomes described in these structural studies were assembled *in vitro* in nuclear extracts, it is possible that CASC3 present in extracts can enter pre-assembled spliceosomes and interact with EJC. Consistently, in the human spliceosome C* structure one of the two CASC3 binding surfaces on EIF4A3 is exposed and available for CASC3 interaction. Still, a possibility remains that, at least on some RNAs or exon junctions, CASC3 assembly may occur soon after splicing within peripseckles as previously suggested (Daguenet et al., 2012).

CASC3 is a more prominent component of cytoplasmic EJCs within mRNPs that have not yet been translated or are undergoing their first round of translation (Figure 5). Previously described functions of CASC3 within translationally repressed neuronal transport granules (Macchi et al., 2003) and posterior-pole localized *oskar* mRNPs in *Drosophila* oocytes (van Eeden et al., 2001) further support CASC3 being a component of cytoplasmic pre-translation mRNPs. It remains to be tested if, like its active role in *oskar* mRNA localization and translation repression (van Eeden et al., 2001), CASC3 also plays a direct role in translational quiescence of RP mRNAs and/or their mobilization to rapidly meet increased demand for translation apparatus (Geyer et al., 1982; Patursky-Polischuk et al., 2009). Nonetheless, our results emphasize the idea of post-transcriptional regulons wherein mRNPs encoding functionally related activities are under coordinated translational control. Capture of such regulons via CASC3-EJC highlights a potential utility of EJC as a molecular marker to identify coordinately translated regulons, perhaps as a complementary approach to the recently described translation-dependent protein knock-off to monitor first round of translation of single mRNPs (Halstead et al., 2015).

### EJC composition and NMD

Based on the binary EJC composition switch, EJC-dependent steps in the NMD pathway can be divided into at least two phases wherein the two alternate factors perform distinct functions. The susceptibility of all tested NMD targets to RNPS1 levels suggests that this protein, and perhaps other components of SR-rich EJCs, serve a critical function in an early phase in the pathway. Such a function could be to recruit and/or activate other EJC/UPF factors either in the nucleus or even during premature translation termination as part of the downstream EJC. Following the compositional switch, the EJC core maintains the ability to activate NMD as it can directly communicate with the NMD machinery via UPF3B (Buchwald et al., 2010; Chamieh et al., 2008). Still, following CASC3 incorporation into the EJC, its ability to stimulate NMD is likely reduced. Such a reduction may stem from the loss of RNPS1 or SR proteins, which are known to enhance NMD (Figure S7, (Aznarez et al., 2018; Sato et al., 2008; Viegas et al., 2007; Zhang and Krainer, 2004)). It is possible that CASC3 overexpression causes the compositional switch to occur at a faster rate or on a greater proportion of mRNAs, or both. Nevertheless, (over)expressed NMD reporters (e.g. β39 mRNA) and some endogenous mRNAs depend on both early and late EJC compositions for their NMD. Notably, the β-globin NMD reporter was previously shown to undergo biphasic decay with faster turnover around the nuclear periphery and slower decay in more distant cytoplasmic regions (Trcek et al., 2013). More recently, singlemolecule imaging of reporter RNPs showed that a fraction of their population diffuses for several minutes and micrometers away from the nucleus before undergoing first round of translation (Halstead et al., 2015). It remains to be seen if mRNPs that are first translated in distant cytoplasmic locales, including those localized to specialized compartments such as neuronal dendrites and growth cones, may rely more on a CASC3-dependent slower phase of NMD. The EJC compositional switch may also underlie the distinct NMD branches identified earlier via tethering of RNPS1 and CASC3 to reporter mRNAs (Gehring et al., 2005). Consistent with these observations, we find that RNPS1 co-purifies more strongly with UPF2, while CASC3 appears to interact more with UPF3B (Figure 2 and data not shown). The nature of the relationship between EJC composition and UPF2 and UPF3B-independent NMD branches is an important avenue for the future work.

## ACKNOWLEDGEMENTS

We would like to thank Pearlly Yan and the OSU Comprehensive Cancer Center genomics core for high-throughput sequencing, Philip Sharp and Paul Boutz for detained-intron datasets, Daniel Schoenberg for H1299 cell line Ribo-Seq and RNA-Seq datasets for translation efficiency estimation, Jens Lykke-Andersen for plasmids and antibodies, Akila Mayeda for anti-RNPS1 antibody, Can Cenik for advice, and Anita Hopper for comments on the manuscript. This work was supported in part by an allocation from the Ohio Supercomputer Center. Funding for this work was provided by the Ohio State University and National Institutes of Health (R01-GM120209) to G.S.

## AUTHOR CONTRIBUTIONS

Conceptualization, G.S. and R.B.; Investigation, J.W.M., L.A.W., R.P., Z.Y., M.J., and G.S.; Writing - Original Draft, G.S., J.W.M. and L.A.W.; Writing - Review & Editing, G.S., J.W.M., L.A.W., R.P., Z.Y., M.J., V.W. and R.B.; Funding Acquisition, G.S.; Resources, G.S., V.W. and R.B.; Supervision, G.S., V.W. and R.B.

## DATA AVAILABILITY

All high-throughput DNA sequencing data are being submitted to GEO and will be available upon publication under Bioproject PRJNA471492.

